# Power spectrum slope and motor function recovery after focal cerebral ischemia in the rat

**DOI:** 10.1101/242388

**Authors:** Susan Leemburg, Bo Gao, Ertugrul Cam, Johannes Sarnthein, Claudio L. Bassetti

## Abstract

EEG changes across vigilance states have been observed after ischemic stroke in patients and experimental stroke models, but their relation to functional recovery remains unclear. Here, we evaluate motor function, as measured by single pellet reaching (SPR), as well as local EEG changes in NREM, REM and wakefulness during a 30-day recovery period after middle cerebral artery occlusion (MCAO) or sham surgery in rats. Small cortical infarcts resulted in poor SPR performance and induced widespread changes in EEG spectra in the ipsilesional hemisphere in all vigilance states, without causing major changes in sleep-wake architecture. Ipsilesional 1–4 Hz power was increased after stroke, whereas power in higher frequencies was reduced, resulting in a steeper slope of the power spectrum. Multielectrode array analysis of ipsilesional M1 showed that these spectral changes were present on the microelectrode level throughout M1 and were not related to increased synchronization between electrodes. Spectrum slope was significantly correlated with post-stroke motor function.

## Introduction

Sleep disorders frequently occur in stroke patients, and are linked to poorer recovery of motor and cognitive functions [1,2]. As sleep plays an important role in learning and memory formation [3–6] and is required for the motor skill acquisition related neuronal plasticity in healthy animals and humans [7–10], it might fulfill a similar beneficial role in promoting plasticity required for recovery of motor function after stroke [11,12]. In stroke patients, recovery of cognitive function is related to sleep amount and efficiency [13] and acquisition of motor tasks is sleep dependent in stroke patients [14–16]. Moreover, in a rat model of focal cerebral ischemia, disruption of sleep has detrimental effects on infarct size and recovery of sensorimotor function [17,18], while administration of the sleep inducing drugs improved functional recovery [19]. Therefore, strategies to improve long-term functional outcome by targeting sleep might be effective and sleep-specific changes in neuronal activity could be a target of particular interest.

In addition to disrupting sleep and wake patterns [1], stroke alters EEG activity within these states in both patients and animal models; leading to increased theta (6–8Hz) and delta (1–4Hz) power and reduced beta power [20,21]. Furthermore, reduced spindle activity during NREM sleep has been reported in stroke patients and was related to poor outcome [22]. Since slow wave activity (SWA, EEG power between 1–4Hz) during non-rapid eye movement sleep (NREM) has been linked to neuronal plasticity and improvements in motor function in the healthy brain [23,24], NREM SWA may have a similar relation to functional recovery after ischemic injury. Indeed, synchronized slow activity around an ischemic lesion was related to the amount of axonal sprouting [25]. However, it is still unclear if and how changes in NREM slow wave activity after stroke are related to increased plasticity leading to improved behavioral outcomes. Alternatively, increased SWA after stroke may be the result of altered synchronization within the damaged area and/or of reduced innervation due to severed inputs, rather than of neuroplasticity. Moreover, increased post-stroke SWA could simply be a result of disturbed sleep-wake patterns.

In this paper, we investigate the effects of small unilateral somatosensory-cortical infarcts on motor function, sleep-wake architecture and EEG power spectrum in ipsilesional and contralesional intact motor cortex. Although the motor cortex was not structurally damaged, severely disrupted activity and extensive remodeling have been shown to occur during recovery [26]. Altered motorcortical neuronal activity is therefore likely the result of neuronal plasticity, rather than loss of cells. Moreover, the small infarcts in this study did not significantly affect sleep-wake architecture, allowing us to assess the relation between motor function and spectral changes in different vigilance states without the confounding effects of e.g. altered sleep pressure.

We hypothesize that, if NREM sleep indeed plays a unique role in mediating recovery, the post-stroke EEG will show sleep-specific changes (e.g. increased SWA) and that these changes will be related to motor function recovery. These sleep-specific changes would be detected in all recorded motorcortical subregions, without necessarily affecting synchronization between them, even when the amount of NREM sleep is not affected. If instead altered sleep EEG is mainly the result of altered innervation of the recorded cortical areas, or of hyper-synchronization within cortical areas, similar changes in neuronal activity will likely be observed outside of NREM sleep as well. Moreover, hyper-synchronization would result in higher similarity of neuronal activity recorded in different motorcortical subregions. Alternatively, altered EEG could be the result of changes in the inhibition-excitation balance within the cortex without necessarily affecting synchrony between subregions.

To investigate these options, we recorded fine motor function and EEG or surface multielectrode activity during a 30-day recovery period after distal middle cerebral artery occlusion.

## Materials and Methods

### Experimental setup

15 rats were implanted with EEG and EMG electrodes after 3 weeks of training on a single pellet reaching task (SPR). After electrode implantation, they were allowed to recover for 8–9 days, during which SPR training continued. 24-hour baseline EEG recordings were then performed, as well as baseline reaching tests. Focal cortical ischemia by middle cerebral artery occlusion (MCAO) was then induced in 8 rats, the hemisphere contralateral to the rat’s preferred reaching paw. 7 sham-operated rats served as controls. Post-surgery EEG recordings were performed on day 1, 4, 7, 10, and 30. The first post-surgery EEG recording (D1) was performed approximately 20h after surgery. Post-surgery SPR testing was performed on days 2, 5, 8, 11, and 31. Rats were sacrificed after the final SPR session.

6 more rats were similarly trained to perform the SPR task, after which thin-film multi electrode arrays (MEAs) were implanted contralateral to the rat’s preferred paw. Rats were allowed to recover for 8–10 days after electrode implantation; SPR training continued during this time. Then, baseline MEA recordings of the first 3 hours of the light and dark period were performed, after which MCAO was induced as described above. Post-MCAO recordings were performed approximately 20h after MCAO (D1) and on day 4. Post-stroke SPR testing was performed on day 2 and 5.

7 rats were used to assess histological damage in M1 after stroke. MCAO was induced (3 left, 4 right hemispheres) and animals were sacrificed 4 days after stroke.

### Animals

28 male Sprague-Dawley rats (21 MCAO, 7 sham, Harlan), weighing 180–200 g at the start of the experiment, were used. Rats used in the EEG and MEA experiments lived in individual cages on a 12h:12 h light-dark cycle. Rats used in the histological experiment were housed in groups of four. All rats were kept on a feeding schedule throughout the duration of the experiment and received 20–25 g of chow per day. Neither the feeding schedule, nor the surgeries caused significant weight loss. Chow was presented once per day, either after the single pellet reaching task, or at lights-on. On SPR testing or training days, rats additionally received chocolate flavored dustless precision pellets (45 mg, product nr. F0299, Bioserv Inc.). The number of pellets a rat received depended on his reaching success, but did not exceed 50. Water was available *ad libitum* throughout the experiment. All experiments were carried with governmental approval (Licenses 167/2005 and 190/2008, 217/15, Kantonales Veterinäramt Zürich, Switzerland) according to local regulations for the care and use of laboratory animals.

### Single pellet reaching (SPR)

Fine motor function was assessed using a skilled reaching task (SPR), in which a rat retrieved a small food pellet located outside of the testing cage by reaching for it using one forepaw, as described previously [18,27]. The rectangular plexiglass SPR testing cage had a shelve mounted 3 cm above the cage floor on the outside of each short wall, which could be reached by the rats via a 1.4 cm wide window in the cage wall. Small indentations in the shelves aligned with the edges of the window assured constant placement of the pellets (1.5 cm from the inside of the cage).

The first week of three weeks of SPR training consisted of daily 10-minute training sessions, in which pellets were initially placed within easy reach. After the rat reliably consumed pellets, they were moved progressively farther away to encourage use of the forepaw. Once a rat demonstrated paw-preference by making >50% of its reaching attempts in a session with a single paw, pellets were placed in the indentation that could only be reached with that preferred paw. In the next two weeks of training, sessions lasted 15 minutes or until the rat completed 50 reaching attempts. During these sessions, pellets were presented on alternating sides of the reaching box, so that the rat fully repositioned its body prior to a new reaching attempt.

During baseline and post-surgery reaching sessions, pellets were presented on one side of the training cage only, after the rat had moved to the other side. These sessions were videotaped for later verification of the scores. SPR attempts were classified as either successful or failed. In a successful attempt, the rat was able to grasp the pellet and eat it. In a failed attempt, the rat knocked the pellet off the shelf or dropped it. SPR success was calculated as the percentage of successfully obtained pellets out of 50 possible attempts.

### Electrode implantation and MCAO surgery

Rats were anaesthetized using 2–2.5% isoflurane in 30% oxygen and 70% N_2_O for all surgical procedures. Rectal temperature was maintained at 36°C. For rats in the EEG experiments, gold-plated miniature screws serving as epidural electrodes were implanted in the skull over the motor cortex of each hemisphere (rostral electrode: bregma +2 mm, 2 mm lateral; caudal electrode: bregma -2 mm, 2.5 mm lateral), where the caudal electrode served as reference for the rostral one in each hemisphere. Thus, the recorded area included the forelimb representation in the primary motor cortex, but not the infracted area located in the somatosensory cortex (Fig. 1A). Two gold wires inserted in the neck muscles served as EMG electrodes. For the MEA experiments, 2 × 2 mm 32-channel thin-film multielectrode arrays (MEA, FlexMEA36-Om, MultiChannel Systems) were implanted subdurally in the hemisphere contralateral to the rats preferred paw. A 5mm long and 4 mm wide craniotomy was made parallel to midline ranging from 3 mm rostral to 2mm caudal of bregma. A 2 × 2 mm window was then carefully cut in the dura mater, to allow for placement of the MEA on the brain surface. The anteromedial corner of the array was placed 1.5 mm rostral and 1 mm lateral of bregma, with the medial edge of the array parallel to midline. The electrode grid covered the area +1 to -1 mm rostral of bregma and 1.5 to 2.5 mm lateral of bregma. The MEA and craniotomy were covered and fixed in place using with Venus Flow dental cement (Kulzer). The MEA and connector were further secured using Paladur cement (Kulzer) and 3 stainless steel miniature anchor screws inserted in the contralateral skull. MEAs were implanted in the ipsilesional hemisphere, contralaterally to the rat’s preferred paw.

**Figure 1.**
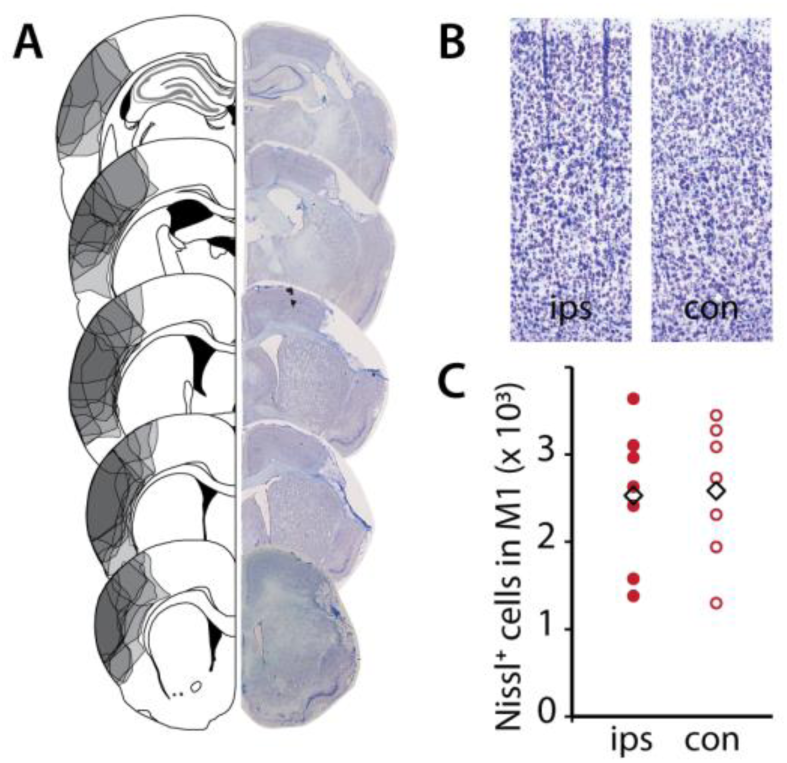
A: Location and extent of ischemic lesions after distal middle cerebral artery occlusion (MCAO). Outlines show lesions for individual animals (N = 8, left), photomicrographs (right) show the infract of a representative animal. B: Representative examples of Nissl-stained primary motor cortex in ipsilesional (ips) and contralesional (con) hemisphere 4 days post-MCAO. C: Cell counts in ipsilesional and contralesional primary motor cortex 4 days post-MCAO. Diamonds indicate mean cell counts in ipsilesional (ips) and contralesional (con) hemisphere. Values for individual animals (N = 7) are shown as circles.

Middle cerebral artery occlusion (MCAO) was performed by electrocoagulating the distal branch of the middle cerebral artery (MCA), using a method adapted from Tamura et al. [28]. The skull overlying the MCA was exposed through an incision in the temporal muscle. A 4-by-5mm window was made in the frontal bone, exposing the underlying vessels and brain. After carefully removing the dura mater covering the artery, the MCA and its main branches were permanently closed using bipolar electrocoagulation. The temporal muscle and overlying skin were then sutured back in place. Sham operated rats underwent the same procedure, but without removal of the dura mater and coagulation of the MCA. All rats were lesioned contralaterally to their preferred paw.

### EEG recording

EEG and EMG activity were recorded using an Embla A10 amplifier and Somnologica Science software (Embla) at 100 Hz and 200 Hz sampling rates, respectively. The EEG signal was filtered using a low-cut filter at 0.3 Hz. EMG was filtered for 50 Hz artifact, and had a low-cut filter at 10 Hz.

EEG was scored as wake, NREM or REM in 8 second epochs based on the contralesional EEG signal and on the EMG using Somnologica Science software (Embla). Wherever the contralesional signal contained many artifacts, the signal from the ipsilesional hemisphere was consulted to help identify the vigilance state. Throughout the experiment, wake was characterized by a low amplitude, high frequency EEG pattern and high EMG activity. NREM was characterized by the occurrence of high amplitude slow waves and tonic low EMG activity. The low amplitude EEG activity of REM was dominated by theta activity and only occasional twitches were present in the EMG signal. Epochs were classified as belonging to a vigilance state when more than half of the epoch fulfilled the criteria for that state. A total period of 22 hours was analyzed for each recording day, starting from 1 hour after the start of the recording at lights-on. Epochs that included artifacts were excluded from spectral analysis.

### MEA recording

Multi-electrode array neuronal activity was recorded using a WS-32 wireless recording system and MCRack software (MultiChannel Systems) at 5 kHz sampling rate with a 1 Hz to 5 kHz signal bandwidth. Signals were low-pass filtered to remove 50Hz artifact, down-sampled to a 100 Hz sampling rate to match the EEG recordings, and divided into 8 second epochs prior to analysis. Signal quality for each channel was assessed based on signal amplitude, waveform and power spectrum. Bad channels, as well as epochs containing movement- or signal-transmission artifacts were excluded from analysis.

On recording days, rats were connected to the wireless system at lights-on and light-off. Due to limited battery life of the wireless transmitters, recordings were limited to a duration of 3 hours, starting 1 hour after the animal was connected at the start of the light or dark period. Animals were habituated to the handling involved in the connecting procedure for at least a week prior to electrode implantation and during the post-implantation recovery period, assuring minimal handling-related disturbance of the animal.

### Data analysis

Total time spent in wake, NREM and REM was calculated for each recording day, as well as the mean episode duration for these vigilance states, was calculated to assess possible stroke-related alterations in sleep-wake patterns. Wake and NREM episodes were defined as periods of ≥24 s preceded and followed by ≥16 s of another vigilance state. REM episodes were defined as being ≥16 s long.

Power spectra for frequencies from 1 to 25 Hz were calculated for artifact-free 8 s epochs for each vigilance state using Welch’s power spectral density estimate (Hamming window, 50% overlap, Matlab 2015, The Math Works, Inc.). Time effects in EEG power spectra were analyzed separately for each experimental condition by performing one-sample t-tests comparing each frequency bin to a 100% baseline value, in order to identify frequency ranges of interest. A bin was considered when the corrected p < 0.05 (corrected for 4 repeated comparisons on day 1, 4, 10 and 30). Total spectral power in delta (1–4 Hz) and beta (15–25 Hz) bands were also calculated for EEG and MEA recordings. EEG spectrum analyses could not be performed for one MCAO animal in wake due to a large number of artifacts.

Spectral slope was calculated as described previously [29]. First, raw power spectra of EEG and MEA recordings were log-transformed by calculating natural logarithm of power values for each frequency bin, after which a linear curve was fitted to the intransformed data to estimate the spectral slope. Relations between EEG power spectrum, spectrum slope and SPR performance were analyzed using repeated measures correlation [30]. Correlations were considered significant when p<0.05.

To assess spatial complexity of the recorded MEA activity as an indicator of altered synchronization between channels, principal component analysis was performed on each artifact-free 8-second epoch of the MEA recording. Detected components were sorted by size and the mean percentage of signal variability explained by the two largest detected components was calculated for each animal. If spatial complexity of MEA activity is low, the amount of variability explained by these components will be high.

Effects of treatment (stroke or sham surgery) and time on SPR performance, sleep-wake architecture and spectrum slope were calculated using repeated measures ANOVA (between-subject factors: treatment (group/ hemisphere), within-subject factor: time). Baseline values for time spent in a vigilance state, episode duration and spectrum slope were included as a covariate to control for possible baseline differences. Statistical analyses were performed using Matlab 2015, SPSS 23 (IBM), and R3.3.3. Results are reported as mean ± s.e.m. and were considered statistically significant when p < 0.05 unless indicated otherwise.

### Histology

Rats were killed with an overdose of sodium pentobarbital and perfused transcardially with 0.1M phosphate-buffered saline (PBS) followed by 4% paraformaldehyde (PFA) in 0.1M PBS. Brains were removed, post-fixated in 4% PFA in 0.1M PBS and cryoprotected in 30% sucrose in 0.1M PBS at 4°C. 30μm thick coronal sections were collected and stained using cresyl violet. Every 12th section was stained and analyzed. Chemicals were purchased from Sigma-Aldrich.

Histological results were analyzed using ImageJ software (U.S. National Institutes of Health). Infarcts were outlined based on the absence of Nissl-stained neurons in the cortex. Infarct volume was then calculated as follows:

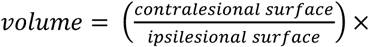
distance between sections.

To assess possible neuronal loss in primary motor cortex (M1) after stroke, M1 was outlined in the ipsilesional and contralesional hemisphere [31]. The number of Nissl+ cell bodies was estimated using the ImageJ particle analyzer plugin, based on micrographs taken at 5x magnification with a Zeiss Axio Scan Z1 microscope.

## Results

### Histological results

Analysis of brain sections showed that middle cerebral artery occlusion (MCAO) resulted in infarcts located in the somatosensory cortex (Fig. 1A). Ischemic damage did not extend into subcortical regions, nor into the motor cortex. Infarct volume in animals in the EEG experiments was 23.9 ± 3.1 mm^3^ (N = 8, Fig. 1A); infarcts in the MEA experiments were slightly smaller: 17.4 ± 1.7 mm3 (N = 6).

MCAO-related somatosensory cortical lesions did not result in significant neuronal loss in primary motor cortex. Estimated numbers of neurons were not significantly different between ipsilesional and contralesional hemispheres (2542 ± 702 ipsilesionally vs. 2597 ± 318 contralesionally, one sided paired-samples t-test, t(6) = -0.750, p = 0.24, N = 7). (Fig. 1B, C)

### Post-stroke motor function impairment and recovery

Motor function was impaired after experimental stroke, even though primary motor cortex did not show histological damage. SPR success was markedly reduced, dropping by on average 60% from baseline performance in MCAO animals (Fig. 2; repeated measures ANOVA, group effect: F(1) = 55.14, p < 0.001; time effect: F(2.7) = 11.83, p < 0.001; time*group interaction F(2.7) = 11.11, p < 0.001). Performance slowly improved and returned to baseline levels on day 31. The acute post-stroke reduction of reaching success on day 2 was significantly correlated to infarct volume (Pearson’s R = -0.713, p = 0.036). However, the amount of subsequent recovery between day 2 and 31 was not (R = 0.384, p = 0.198). Sham surgery had no effect on reaching success.

**Figure 2.**
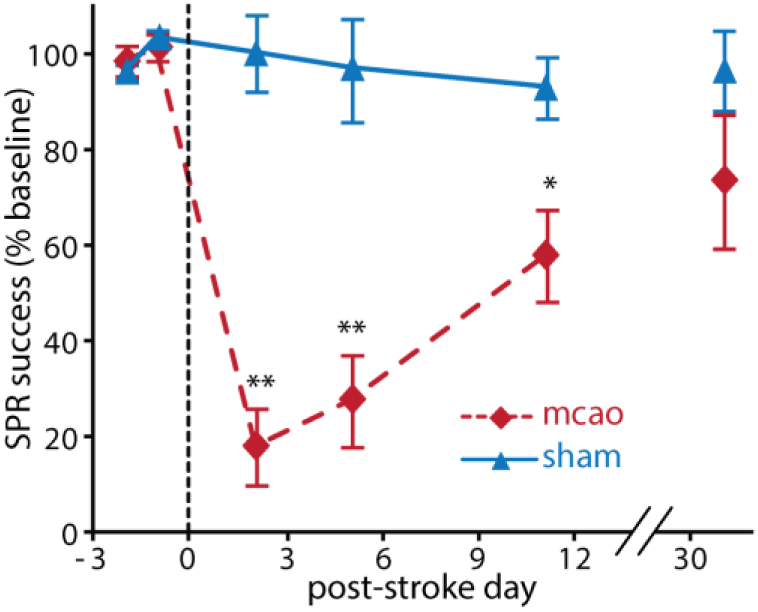
Pre- and post-stroke single pellet reaching performance in MCAO and sham operated animals. Asterisks denote significant differences between groups (* p < 0.05, **p < 0.01, MCAO: N = 8, sham: N = 7)

### Effects of stroke on sleep-wake architecture

The small cortical infarcts did not result in any major stroke-related changes in the amount of time spend awake or in NREM sleep in MCAO rats. The amount of NREM sleep was not significantly different between MCAO and sham-operated rats (repeated measures ANOVA, group effect: F(1) = 1.025, p = 0.331; time effect: F(4) = 2.983, p = 0.028; group*time interaction F(4) = 3.659, p = 0.011), nor was the amount of wakefulness (repeated measures ANOVA, group effect: F(1) = 0.197, p = 0.665; time effect: F(1.856) = 1.042, p = 0.395; group*time interaction F(1.856) = 1.491, p = 0.246) (Table 1). Time spent in REM during the light phase was acutely reduced by ≈35% after stroke, but total time spent in REM sleep was not significantly different in MCAO and sham-operated animals (repeated measures ANOVA, group effect: F(1) = 1.040, p = 0.328; time effect: F(4) = 8.828, p < 0.001; group*time interaction F(4) = 3.165, p = 0.022). Vigilance state consolidation, as reflected by episode duration (Table 2), remained unaltered in MCAO and sham-operated animals. Wake episode duration (repeated measures ANOVA, group effect: F(1) = 4.567, p = 0.054; time effect: F(1.899) = 2.250, p = 0.130; group*time interaction: F(1.899) = 1.658, p = 0.213), NREM episode duration (repeated measures ANOVA, group effect: F(1) = 1.680, p = 0.221; time effect: F(4) = 2.591, p = 0.049; group*time interaction: F(4) = 7.530, p < 0.001) and REM episode duration (repeated measures ANOVA, group effect: F(1) = 0.001, p = 0.972; time effect: F(4) = 2.341, p = 0.068; group*time interaction: F(4) = 0.734, p = 0.573) were not significantly altered by stroke.

### Effects of stroke on EEG power spectra

Although MCAO did not have a major effect on the amount of NREM, REM and wake, EEG activity within each of these states was altered by the infarct. The ipsilesional hemisphere was most obviously affected, resulting in a marked asymmetry in the EEG (Fig. 3). Power spectra for all MCAO animals revealed widespread stroke-related changes in all three vigilance states. EEG was similarly affected in wakefulness (fig. 4A), NREM (fig. 4B) and REM sleep (fig. 4C): a large, highly variable increase in delta power (1–4Hz) was found in all three vigilance states. Additionally, spectral power in all higher frequencies (5–25 Hz) was acutely reduced. In wake and REM, 5–25Hz power returned to baseline levels over the course of recovery. In NREM, however, power in this frequency range did not fully recover to baseline levels. Contralesional power spectra (Fig. S1) in MCAO rats and contralateral and ipsilateral power spectra in sham operated rats (Fig. S2) were unaffected by stroke or surgery. Thus, although the observed spectral changes were hemisphere- and stroke-specific, they were not restricted to a specific vigilance state.

**Figure 3.**
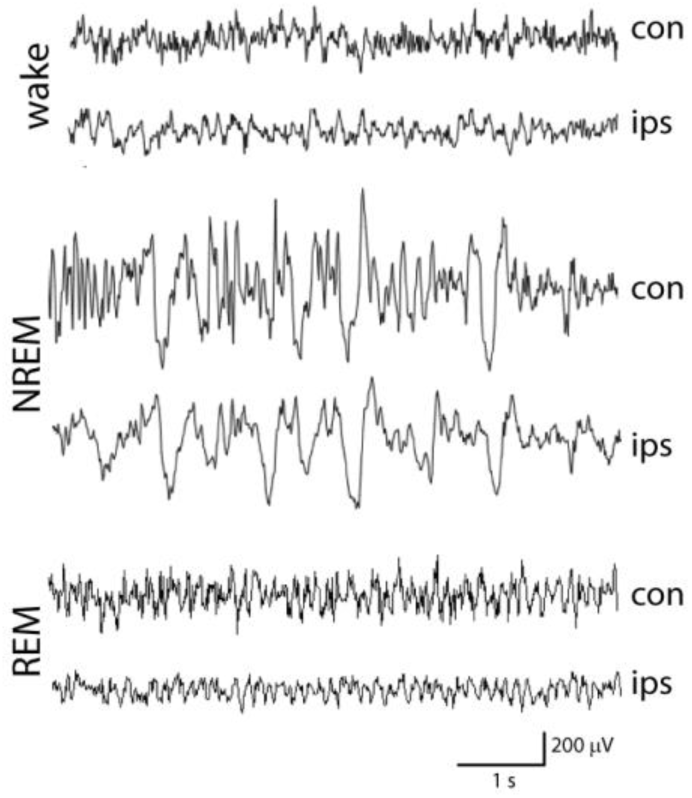
Representative EEG traces of wake, REM and NREM sleep in ipsilesional (ips) and contralesional (con) hemisphere 4 days post-stroke.

**Figure 4.**
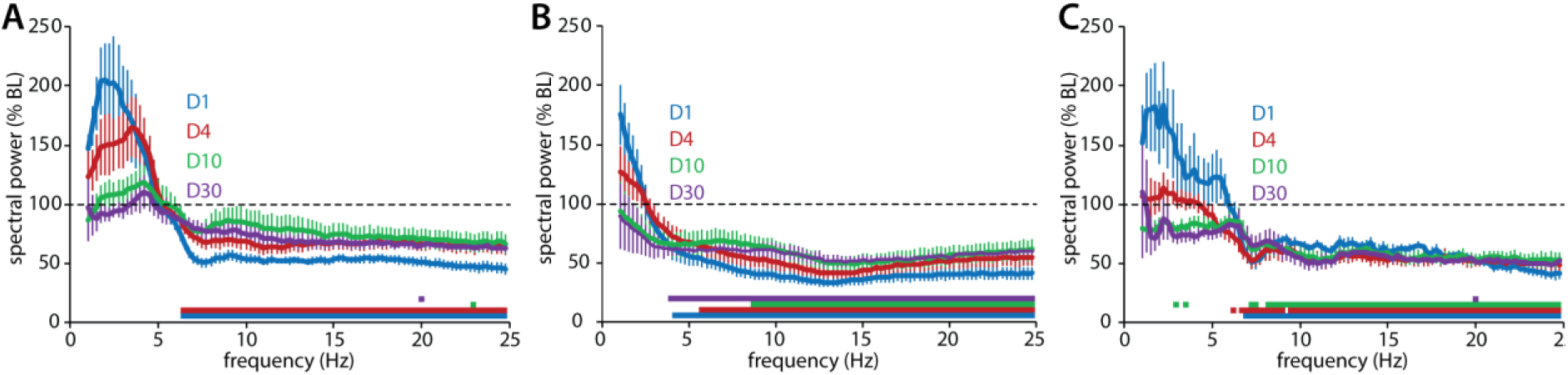
A: Mean ipsilesional wake EEG spectrum change (power as % of baseline) on day 1,4,10 and 30 post-MCAO (N = 7, values are mean ± s.e.m). Squares below the curves indicate significant differences from baseline per frequency bin (p < 0.05). Inset shows power in delta (1–4Hz) and beta (15–25Hz) as percentage of baseline (asterisks indicate significant differences from baseline). B: Mean ipsilesional NREM EEG spectrum change 1,4,10 and 30 days post-MCAO (N = 8, values are mean ± s.e.m). Squares below the curves indicate significant difference from baseline per frequency bin (p< 0.05). Inset shows power in delta (1 -4Hz) and beta (15–25Hz) as percentage of baseline (asterisks indicate significant differences from baseline). C: Mean ipsilesional REM EEG spectrum change 1, 4, 10 and 30 days post-MCAO (N = 8, values are mean ± s.e.m). Squares below the curves indicate significant difference from baseline per frequency bin (p < 0.05). Inset shows power in delta (1 -4Hz) and beta (15–25Hz) as percentage of baseline (circles and asterisks indicate significant differences from baseline).

Because stroke affected the entire frequency range of the EEG power spectrum, we have opted to quantify the effects of stroke on the shape of the entire spectrum, rather than to discuss the effects on traditional frequency bands. To this end, we estimated the linear slope of the ln-transformed EEG power spectrum (fig. 5A). The combination of increased low frequency activity and reduced higher frequency EEG activity resulted in altered slopes of post-stroke power spectra. As a result, spectrum slopes were significantly steeper than baseline in wake, NREM and REM in ipsilesional cortex of MCAO rats on post-stroke day 1 and 4, but not in contralesional cortex or in sham-operated animals. Wake spectrum slopes were significantly affected by stroke in (repeated measures ANOVA, treatment effect: F (3) = 6.074, p = 0.003; time effect: F(2.65) = 12.83, p < 0.001; treatment*time interaction F(7.951) = 4.379, p < 0.001, Fig. 5B). Similar changes were found in REM (repeated measures ANOVA, treatment effect: F (3) = 6.133, p = 0.003; time effect: F(2.318) = 7.799, p < 0.001; treatment*time interaction F(6.953) = 5.081, p < 0.001, Fig. 5D), and in NREM sleep (repeated measures ANOVA, treatment effect: F (3) = 3.970, p = 0.020; time effect: F(2.993) = 2.549, p = 0.044; treatment*time interaction F(8.978) = 5.627, p < 0.001, Fig. 5C). Although ipsilesional NREM higher frequency power remained below baseline levels on day 10 and 30, spectrum slope has normalized on these days. This is likely the result of a concurrent decrease in low frequency power, resulting in a baseline-like spectrum shape (i.e. a mostly flat curve in Fig. 4A).

**Figure 5.**
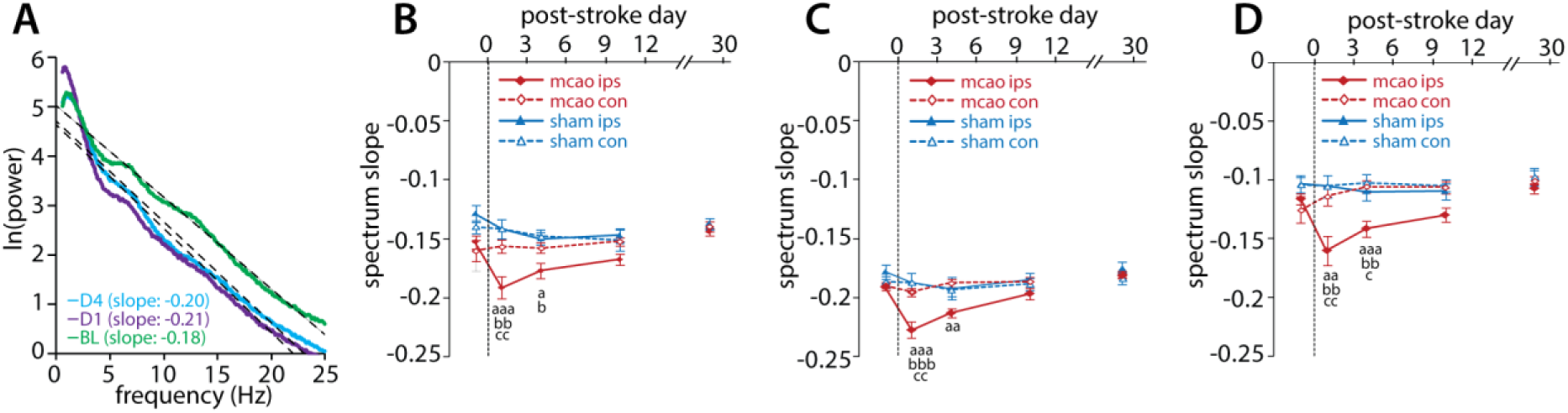
A: Representative NREM spectra with fitted linear curves estimating overall spectrum slope at baseline (BL), 1 and 4 days post-MCAO. B: Time course of wake EEG spectrum slope in ipsilesional (ips) and contralesional(con) hemispheres of MCAO and sham-operated animals. Values are mean ± s.e.m, sham: N = 8, mcao: N = 7. Letters denote significant differences between groups: mcao ips vs. mcao con:^a^ p < 0.05, ^aaa^p ≤ 0.001; mcao ips vs. sham con: ^bb^ p < 0.01; mcao ips vs. sham ips: ^c^ p < 0.05, ^ccc^ p ≤ 0.001. C: Time course of NREM EEG spectrum slope in ipsilesional (ips) and contralesional(con) hemispheres of MCAO and sham-operated animals. Values are mean ± s.e.m, sham: N = 8, mcao: N = 8. Letters denote significant differences between groups: mcao ips vs. mcao con: ^aa^ p ≤ 0.01, ^aaa^p < 0.001, mcao ips vs. sham con: ^bbb^p ≤ 0.001, mcao ips vs. sham ips:^cc^p < 0.01. D: Time course of REM EEG spectrum slope in ipsilesional (ips) and contralesional (con) hemispheres of MCAO and sham-operated animals. Values are mean ± s.e.m, sham: N = 8, mcao: N = 8. Letters denote significant differences between groups: mcao ips vs. mcao con:^aa^ p < 0.01, ^aaa^p ≤ 0.001, mcao ips vs. sham con: ^bb^p < 0.01, mcao ips vs. sham ips: ^c^ p < 0.05,^cc^ p < 0.01.

### Relation between EEG spectrum and behavioral outcome

If the described effects of stroke on power spectra play in a role in mediating recovery of fine motor function after stroke, the degree of stroke-induced EEG change should be related to the amount of functional recovery.

Stroke not only had opposite effects on slow and fast EEG activity, slow and fast EEG activity had opposing relationships to post-stroke motor function as well. Post-stroke SPR performance (% of baseline) was negatively correlated to the post-stroke increase in 1–5 Hz power in wake and REM (% of baseline, Fig. 6A). A similar negative, but non-significant correlation was found in this frequency range in NREM. By contrast, SPR performance was positively correlated to high-frequency activity in REM and wake (20–25 Hz) and in a wide range of frequencies in NREM (8–25 Hz).

**Figure 6.**
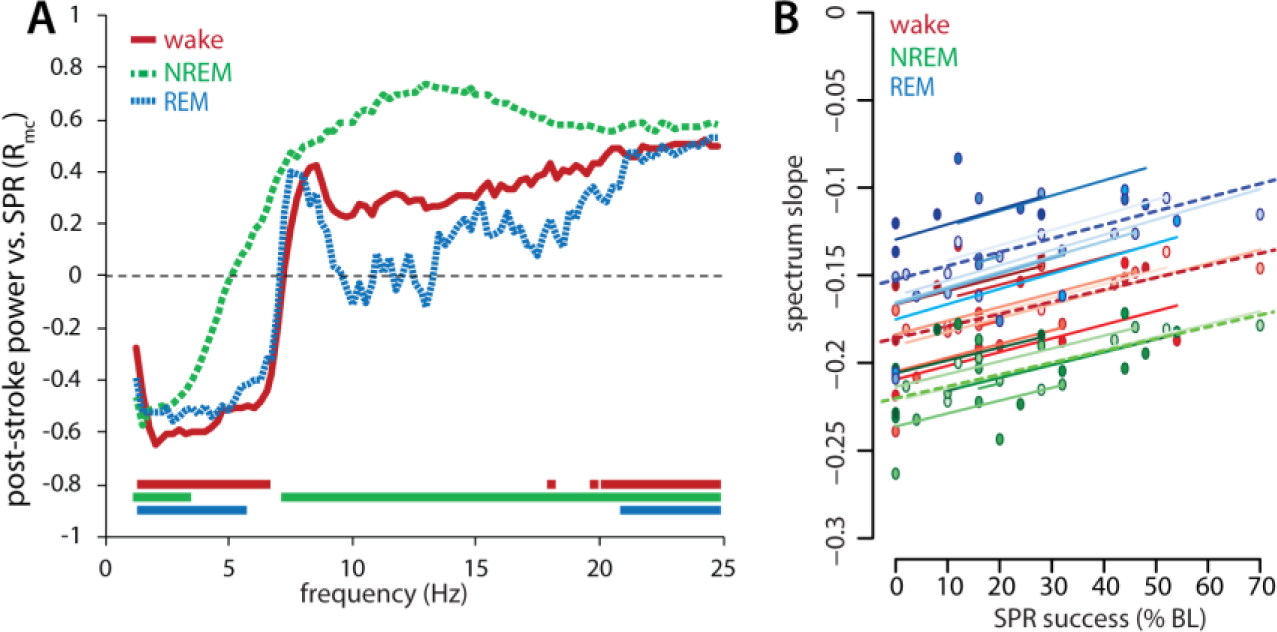
A: Pearson’s R value showing correlation between ipsilesional EEG power spectrum change (% of baseline) and pellet reaching performance (% of baseline) post-stroke in MCAO rats (N=7 for wake, N = 8 for NREM and REM). Squares indicate statistically significant correlations per frequency bin (p < 0.05). B: Repeated measures correlation between post-stroke pellet reaching performance (% of baseline) and ipsilesional power spectrum slope in wake (N = 7), NREM (N = 8) and REM (N = 8). Solid lines indicate individual correlations for each animal, dashed lines indicate overall correlation for wake (red), REM (blue), and NREM (green).

Thus, high levels of low frequency power and low levels of high frequency EEG activity in a recording session predicted poor motor function on the next day. This is also apparent in the relation betweenpost-stroke spectrum slope and motor function: reduced high-frequency activity and increased delta power resulted in abnormally steep power spectrum slopes that were related to poor function (Fig. 6B). Indeed, shallower spectrum slopes were positively correlated to next day motor function in wake (R = 0.649, p < 0.001), REM (R = 0.602, p = 0.001) and NREM (R = 0.625, p < 0.001).

### Spatial changes in local cortical activity after stroke

The MCAO related changes in the EEG could indicate altered synchronization between cortical subregions. Alternatively, steeper spectrum slopes could indicate altered signal-to-noise ratio, or changes in inhibition-excitation balance in the cortex that are not related to local synchronization. To investigate local effects of MCAO within ipsilesional motor cortex, neuronal activity was recorded using 32-channel thin-film multi electrode arrays (MEA). Because the effects of MCAO on EEG power spectra and their relation to motor function were similar in sleep and wakefulness, MEA power spectra were analyzed without differentiating between vigilance states.

Stroke-induced power spectral changes in MEA recordings were similar to those found in EEG and consisted of increased low frequency power and a widespread decrease of power in higher frequencies. Most MEA channels showed increased delta band power on day 1 post-MCAO (Fig. 7A,B). Mean values of all channels showed that delta power had mostly returned to baseline levels by day 4 (repeated measures ANOVA, F(1.04) = 8.796, p = 0.029, Fig. 5B). Similar to EEG, MEA beta band power (15–25 Hz) was acutely reduced on day 1 (Fig. 7A,C). Mean beta power of all channels had recovered by day 4 (repeated measures ANOVA, F(1.07) = 10.18, p = 0.021). Thus, while similar in frequency range, the effects of stroke on MEA power spectra was not as long lasting as those seen in EEG. It is unclear if this is the result of the MEA electrode size and configuration, of the slightly smaller infarcts in MEA-implanted animals, or a combination of both.

**Figure 7.**
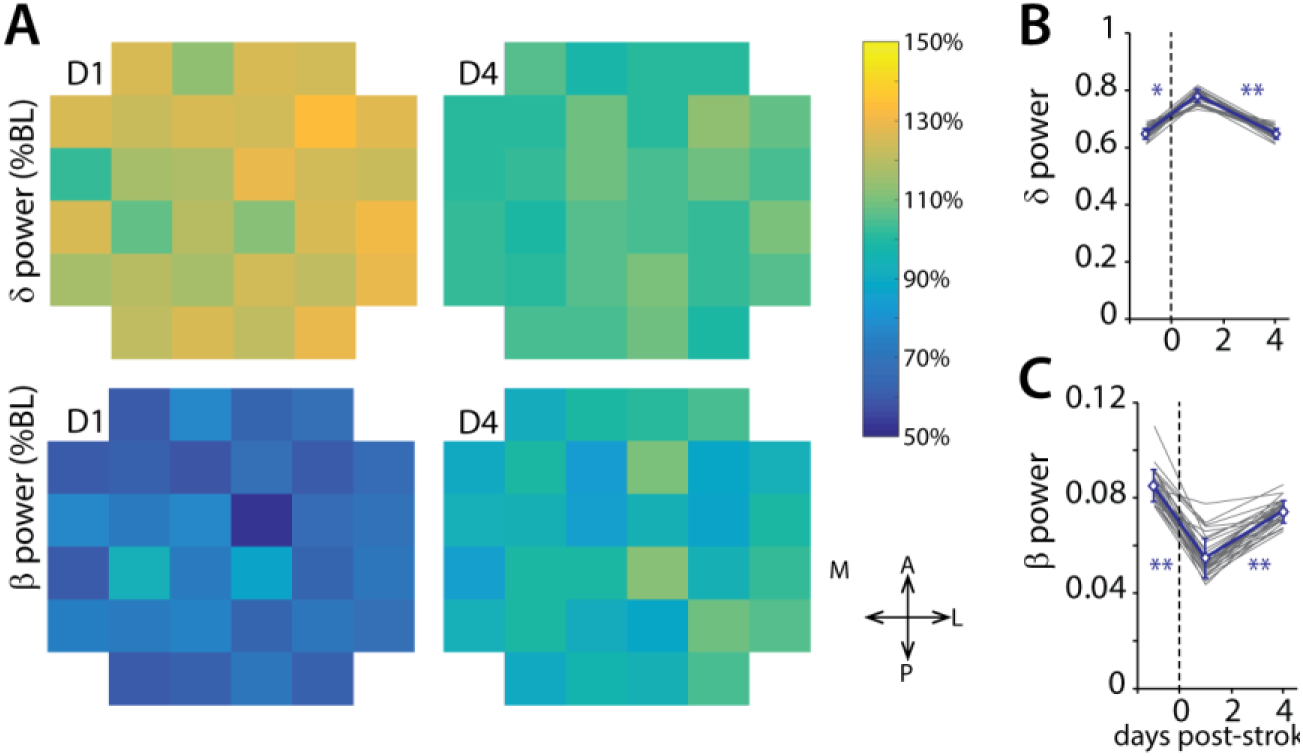
A: Localized changes in beta and delta band power on D1 and D4 post-stroke. Values shown as percentage of baseline per electrode, mean of 6 animals. Arrows denote array orientation (A = anterior, P = posterior, M = medial, L = lateral) B: Mean delta band power at baseline, D1 and D4 post-stroke for individual MEA electrodes (gray) and mean values ± s.e.m for all electrodes (blue). Asterisks denote significant difference between groups (* p < 0.05, ** p < 0.01). C: Mean beta band power at baseline, D1 and D4 post-stroke for individual MEA electrodes (gray) and mean values ± s.e.m for all electrodes (blue). Asterisks denote significant difference between groups (** p < 0.01).

The observed changes in delta and beta power varied across the 32 channels of the multielectrode array, indicating local variability in motorcortical response to a somatosensory cortical lesion. Changes in both delta and beta power were slightly larger in electrodes located in lateral parts of the array, which were located in closer proximity to the infarct (Fig. 7A). As a result, the steepest post-stroke spectrum slopes were found in these channels as well (Fig. 8A). Spectrum steepness was increased in the majority of MEA channels, although the magnitude of this change was variable. As a result, mean spectrum slope changes after MCAO were not statistically significant (repeated measures ANOVA, F(1) = 5.893, p = 0.060, Fig. 8B).

**Figure 8.**
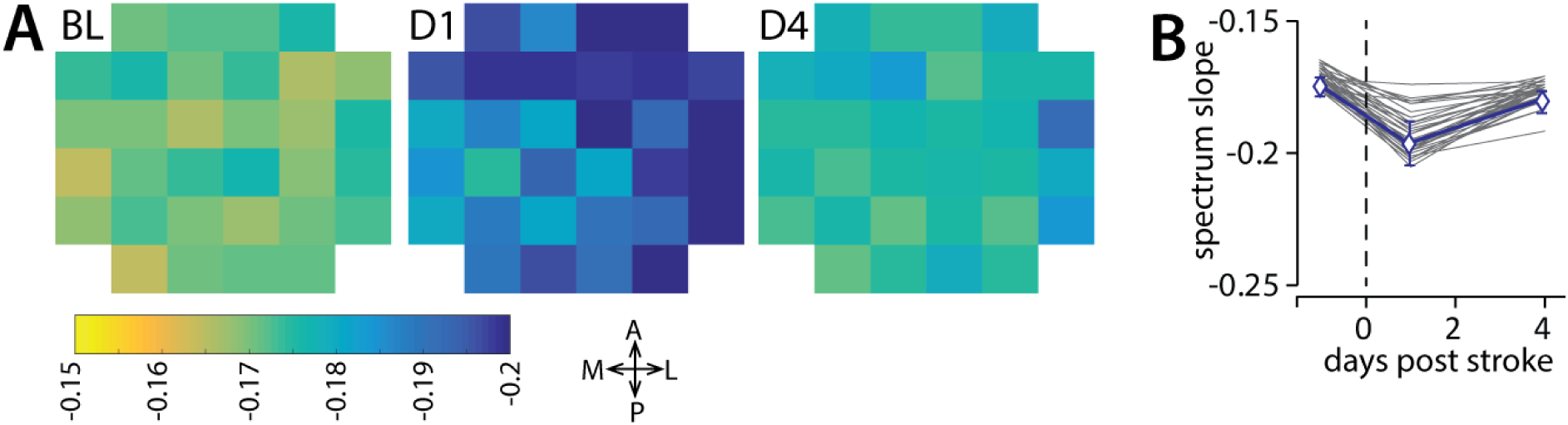
A: Spectrum slopes at baseline, D1 and D4 post-stroke for individual electrodes. B: Mean spectrum slope at baseline, D1 and D4 post-stroke for individual MEA electrodes (gray) and mean values ± s.e.m for all electrodes (blue). N = 6 animals

To investigate possible stroke-related changes in synchronization between cortical subregions, the spatial complexity of the MEA activity was estimated by calculating the percentage of signal variability explained by the largest components detected using principal component analysis (Fig. 9A). Overall spatial complexity of cortical activity in was not significantly affected by stroke (Fig. 9B; repeated measures ANOVA, F(1.073) = 0.290, p = 0.632). Spatial complexity of wake epochs alone, which likely reflects cortical function related to motor task activity more accurately than combined activity in all vigilance states, was also not affected by stroke (repeated measures ANOVA, F(1.030) = 0.593, p = 0.488).

**Figure 9.**
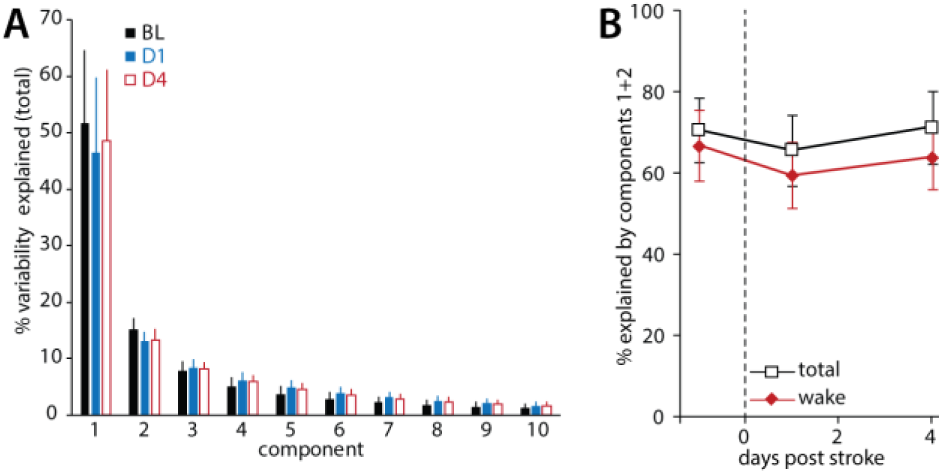
A: Spatial complexity of MEA signals in all vigilance states at baseline (BL), D1 and D4 as reflected in the largest components obtained after principal component analysis of the MEA signal. B: Mean variability explained by components 1 and 2 at baseline, D1 and D4 post-stroke in all vigilance states (total) and in wake. Values shown are mean ± s.e.m, N = 6 animals.

## Discussion

In this paper, we analyzed the long-term effects of focal cortical ischemia on EEG activity in wakefulness, NREM and REM sleep and its relation to motor function recovery in the rat. MCAO led to reduced single pellet reaching performance as well as major changes in ipsilesional power spectra, both of which recovered during the 30-day recovery period.

In contrast with previous studies in rats [17–19], we did not find any major changes in the amount of sleep and wakefulness, nor in vigilance state consolidation. The absence of changes in sleep-wake architecture may be related to the small infarct volumes in our animals compared to previous studies. A small reduction in REM sleep amount was detected in the light-phase on day 1 post-stroke, although the total amount of REM on that day did not change. This is in line with previous studies showing reduced REM sleep acutely after brain injury [18,19]. Possibly, NREM and wake patterns had already normalized by the start of recording, approximately 20 hours after surgery.

Post-stroke changes in ipsilesional EEG power spectra were similar in wake, NREM and REM sleep: power in the slow wave frequency range was increased, whereas activity in higher frequencies was reduced. The changes in slow and fast EEG activity had opposite relations to post-stroke motor function: high levels of low frequency activity were related to poor function, and high levels of high frequency activity instead were correlated to good SPR performance. The shifts in low and high frequency EEG power resulted in acutely steeper slopes of the power spectrum, which recovered along with SPR performance. Although high frequency EEG power does not recover in all states, proportions of low and high frequency power have normalized by the end of the recovery period, resulting in slopes that are similar to baseline spectra. Shallower, more baseline-like slopes were correlated to better post-stroke motor function. These results are similar to those found in patients [20,32–35].

A negative correlation between high levels of delta power and poor motor function seems contrary to its role in memory consolidation, particularly for NREM sleep. If perilesional slow waves are a cue for axonal sprouting [25], the newly formed projections may only contribute to motor function recovery in later stages of recovery [36]. Moreover, high delta activity in wake is linked to poor performance in healthy animals [37]. Hypothetically, it is possible that pharmacologically increasing delta activity specifically in NREM sleep leads to improved recovery in part due to a homeostatic reduction in delta activity in wake and REM sleep, although this remains to be studied. Performing a skilled reaching task in itself has been shown to increase delta activity immediately after training [23]. In the current study, we recorded EEG on the day prior to and approximately 48 hours after SPR sessions. As training effects on slow wave activity dissipate within hours after training, the observed increases in slow wave activity after stroke changes are unlikely to be the result of motor skill testing itself. Moreover, slow wave activity remained unaffected in sham-operated animals.

Although delta and beta activity alone were both related to post-stroke recovery, the slope of the ln-transformed power spectrum takes both into account simultaneously. Both beta and delta activity did not fully return to baseline levels at the end of the 30-day recovery period, although their relative proportions had normalized by then, as did the overall shape of the power spectrum. The spectrum slope reflects the excitation inhibition balance in cortical activity during different behavioral states [38,39]. The changes in spectrum slope were similar in wake, NREM and REM sleep, indicating that, in the absence of sleep disturbances, ipsilesional M1 showed heightened levels of inhibition in all vigilance states. This is consistent with excessive tonic inhibition mediated by extrasynaptic GABA-A receptors in the undamaged perilesional cortex [40]. Steeper slopes were found in all recorded subregions within the ipsilesional M1, but were slightly more pronounced closer to the infarct. Changes in spectrum slope that could indicate altered excitation-inhibition balance, did not lead to increased synchronization of spontaneous neuronal activity between cortical subregions, even though a marked simplification of motor map responses to intracortical microstimulation has been described [26]. It is possible that such changes in spatial complexity of motorcortical activity are most pronounced during task specific behavior, i.e. skilled reaching.

Beneficial effects of stimulants [40,41], as well as sleep-promoting drugs [19] may be linked to a role in normalizing the cortical excitation inhibition balance. Whereas stimulants directly increase excitation, NREM sleep promotion could lead to normalization via synaptic scaling [24]. A such, stimulant- and sleep-promoting drugs might have complementary use in promoting functional recovery, provided their effects remain within the appropriate vigilance state. Spectrum slope could be a useful readout of cortical activity to guide such interventions.

